# Textile waste and microplastic induce activity and development of unique hydrocarbon-degrading marine bacterial communities

**DOI:** 10.1101/2020.02.08.939876

**Authors:** Elsa B. Girard, Melanie Kaliwoda, Wolfgang W. Schmahl, Gert Wörheide, William D. Orsi

## Abstract

Biofilm-forming microbial communities on plastics and textile fibers are of growing interest since they have potential to contribute to disease outbreaks and material biodegradability in the environment. Knowledge on microbial colonization of pollutants in the marine realm is expanding, but metabolic responses during substrate colonization remains poorly understood. Here, we assess the metabolic response in marine microbial communities to three different micropollutants, virgin high-density polyethylene (HDPE) microbeads, polysorbate-20 (Tween), and textile fibers. Intertidal textile fibers, mainly cotton, virgin HDPE, and Tween induced variable levels of microbial growth, respiration, and community assembly in controlled microcosm experiments. RAMAN characterization of the chemical composition of the textile waste fibers and high-throughput DNA sequencing data shows how the increased metabolic stimulation and biodegradation is translated into selection processes ultimately manifested in different communities colonizing the different micropollutant substrates. The composition of the bacterial communities colonizing the substrates were significantly altered by micropollutant substrate type and light conditions. Bacterial taxa, closely related to the well-known hydrocarbonoclastic bacteria *Kordiimonas* spp. and *Alcanivorax* spp., were enriched in the presence of textile-waste. The findings demonstrate an increased metabolic response by marine hydrocarbon-degrading bacterial taxa in the presence of microplastics and textile waste, highlighting their biodegradation potential. The metabolic stimulation by the micropollutants was increased in the presence of light, possibly due to photochemical dissolution of the plastic into smaller bioavailable compounds. Our results suggest that the development and increased activity of these unique microbial communities likely play a role in the bioremediation of the relatively long lived textile and microplastic pollutants in marine habitats.

**Graphical Abstract:** 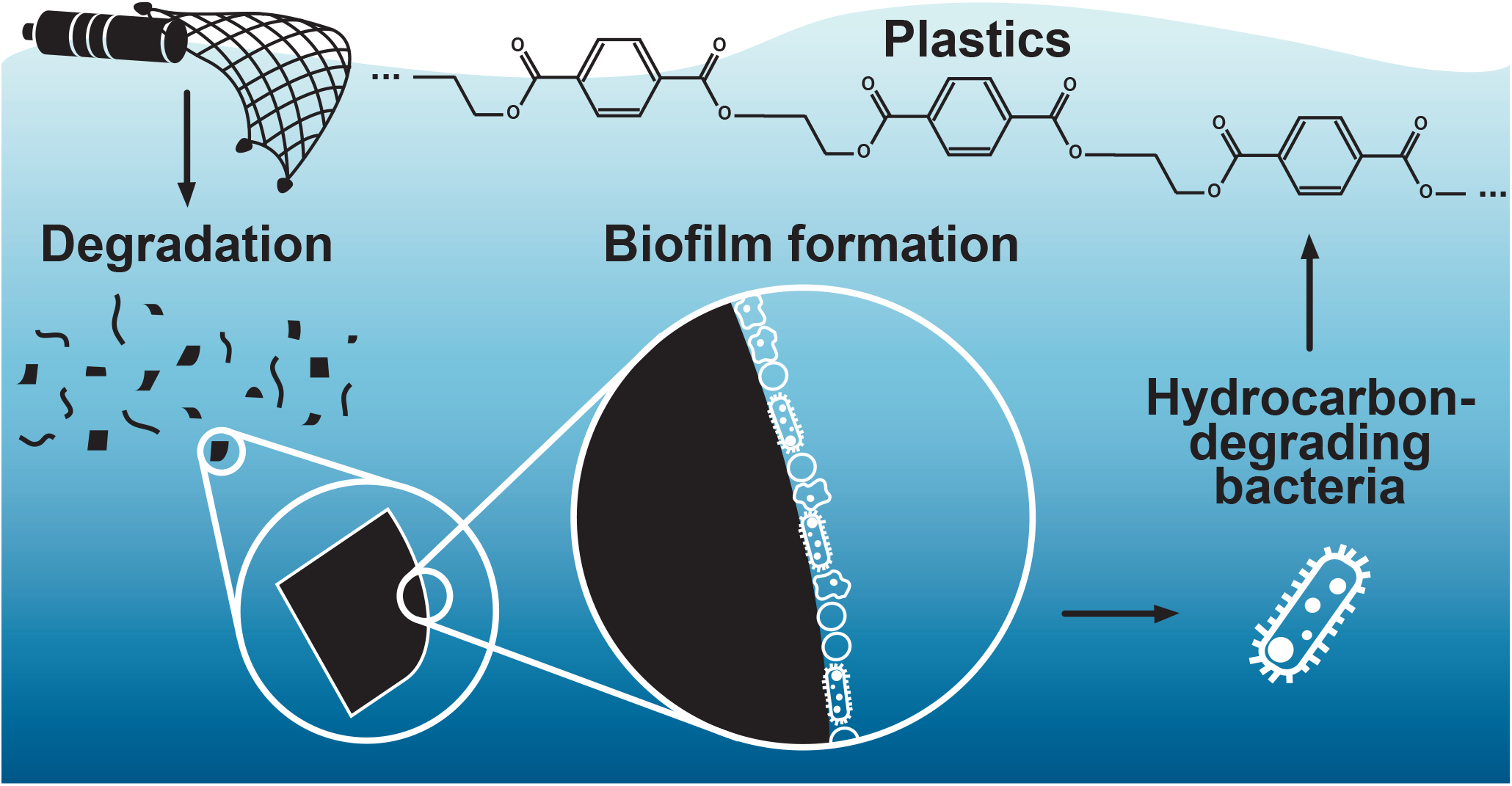

## INTRODUCTION

Plastics are synthetic organic polymers that are composed of long chains of monomers primarily made from petrochemical sources ^1^. The mismanagement of waste in regions with high coastal population density has been linked to high plastic input into the ocean, resulting in an annual flow of 4.8 to 12.7 million tons per year since 2010 ^2^. Once released in the environment, debris are readily colonized by complex microbial communities ^3^. Consequently, macro- and micro-litter may facilitate microbial dispersal throughout the marine realm. However, knowledge gaps regarding the mechanisms of microbial biodegradation of plastic and textile waste.

Plastic-degrading microorganisms have been studied since the 1960’s. Summer studied the inhibition of microorganism growth for lasting polymers, to counter the deterioration of plastics due to mold and bacteria, because some plasticisers (chemical additives used to provide strength and flexibility), e.g., Ester-type plasticisers, are in turn a source of nutrients, which sustains microbial activity leading to natural degradation of the polymer ^4^. More recent studies have shed some light on the diversity of microbial communities colonizing synthetic polymers. For example, Zettler *et al.* (2013) identified a highly diverse microbial community settled on plastic debris, termed as the “plastisphere”, unraveling polymer-dependent communities ^5^. Moreover, its species richness appears to be more important than the microbial community in seawater samples for a given surface ^3,5^. The colonization of plastic debris by bacteria is hypothesized as a two-step settlement: primary colonization by α- and γ-proteobacteria, and subsequent secondary colonization by Bacteroidetes ^6,7^.

Bioremediation of plastic pollution can be aided by heterotrophic bacteria ^8^. These microorganisms may survive by extracting the carbon from plastic particles, via hydrolysis of the hydrocarbon polymer ^7,8^. Recent studies have identified a few bacterial species able to deteriorate plastics, for example, *Ideonella sakainesis*, which is a betaproteobacterium actively degrading polyethylene (PE) ^9^. The plastisphere also harbours a variety of potential pathogens, i.e., harmful microorganisms to animals, such as *Vibrio* spp. ^3,5,6^. Indeed, Lamb et al. observed the transfer of harmful bacteria from plastic litter to reef-building corals, causing three diseases (skeletal eroding band disease, white syndromes and black band disease), which led to coral mortality ^10^. Such observation highlights the need for reducing and taking action on plastic pollution in the environment.

Microparticles of textile waste (i.e., synthetic and natural fibers) enter the ocean due to atmospheric deposition and poor wastewater incubation plant filtration systems allowing the leakage of fibers to the aquatic environment, making it one of the most abundant and recorded micropollutants at sea ^11,12^. Thus, this study aims to assess the potential for bioremediation of microplastics and textile waste by marine microbial communities in a controlled microcosm experiments.

The main questions addressed here are whether specific micropollutant-associated microbial communities develop in the presence of high-density polyethylene (HDPE) microbeads and textile fibers as sole source of carbon, and how these substrates influence their metabolic state. Furthermore, we investigated whether light has an impact on the development and metabolism of these communities. We hypothesized that hydrocarbon-degrading microbes can use microplastics and textile waste as the sole carbon source, and that different types of microplastics will select for unique communities with different levels of activity. Because light also plays a role in the abiotic degradation of organic matter in aquatic environments ^13^, we hypothesized that exposure to light may also improve the ability of the bacteria to utilize carbon from plastic polymers as a growth substrate due to its photochemical dissolution. The results contribute to our understanding of the formation and development of plastics and textile-waste-associated microbes and their potential role in bioremediation of these widespread environmental micropollutants.

## MATERIAL AND METHODS

### Experimental setup and sampling

A total of 15 mL of aquarium seawater containing microbial communities were incubated in 20 mL glass petri dishes for 108 h at room temperature, which received either no micropollutants (control), polysorbate (Tween) 20, Tween 20 and HDPE microbeads, or textile fibers (Tab. 1, Fig. 1). Tween 20 was used as a control since it is used as an emulsifier for the HDPE microbeads and thus serves to test whether the microbes respond only to the Tween or also are effected by the HDPE itself. The artificial seawater microbial community comes from an aquarium (642 L) built of imported live rocks, which hosts many reefs organisms, such as hexacorals, octocorals, gorgonians, sea anemones (Aiptasia sp.), marine sponges (Lendenfeldia chondrodes, Tethya wilhelma), marco-algae (Chaetomorpha linum) and cyanobacteria, mussels (Mytilus edulis) and benthic foraminifera (Elphidium crispum) (Fig. 1A). For each of these incubations, one set was placed under LEDs (Mitras LX6200 HV; light spectrum of 380 nm to 700 nm) with a 12/24 h light cycle (referred to as “light”) and the other one placed inside a cardboard box covered with aluminium foil to block incoming light (referred to as “dark”) (Fig. 1). Each incubation set consists of twelve glass petri dishes sealed with parafilm containing a submerged oxygen sensor spot (PreSens Precision Sensing): three controls and nine incubations (Fig. 1B). The oxygen sensor spot was positioned at the bottom of the petri dish to measure the minimal concentration of O2, that could be dissolved into the bottom water of the petri dish after diffusion from the overlying headspace.

**Table 1.** Description of each incubation type, equally exposed to light and dark conditions.

|  |  |
| --- | --- |
| Control | Control petri dishes containing only 15 mL of artificial seawater, with no micropollutants. |
| Tween | 0.1% Tween 20 solution produced following the solubilisation protocol of Cospheric LLC (https://www.cospheric.com/).<br>150 μL of tween solution was added to 15 mL of artificial seawater.<br>Final tween concentration: 0.01 mg/mL. |
| HDPE | Virgin HDPE microbeads (1-4 μm; 0.96 g/cm<sup>3</sup>) solubilized in 0.1% Tween 20 solution (final concentration of 2.6% solid solution of HDPE).<br>150 μL of HDPE solution was added to 15 mL of artificial seawater.<br>Final HDPE concentration: 0.26 mg/mL. |
| Textile fiber | Ca. 500 textile fibers were collected from intertidal sediment at Coral Eye Resort (Bangka Island, North Sulawesi, Indonesia). Fibers were washed twice in Ethanol >99%.<br>150 μL of Milli-Q H<sub>2</sub>O containing 50-60 fibers was added to 15 mL of artificial seawater.<br>Final fiber concentration: 3.5 fibers/mL. |

**Figure 1.**
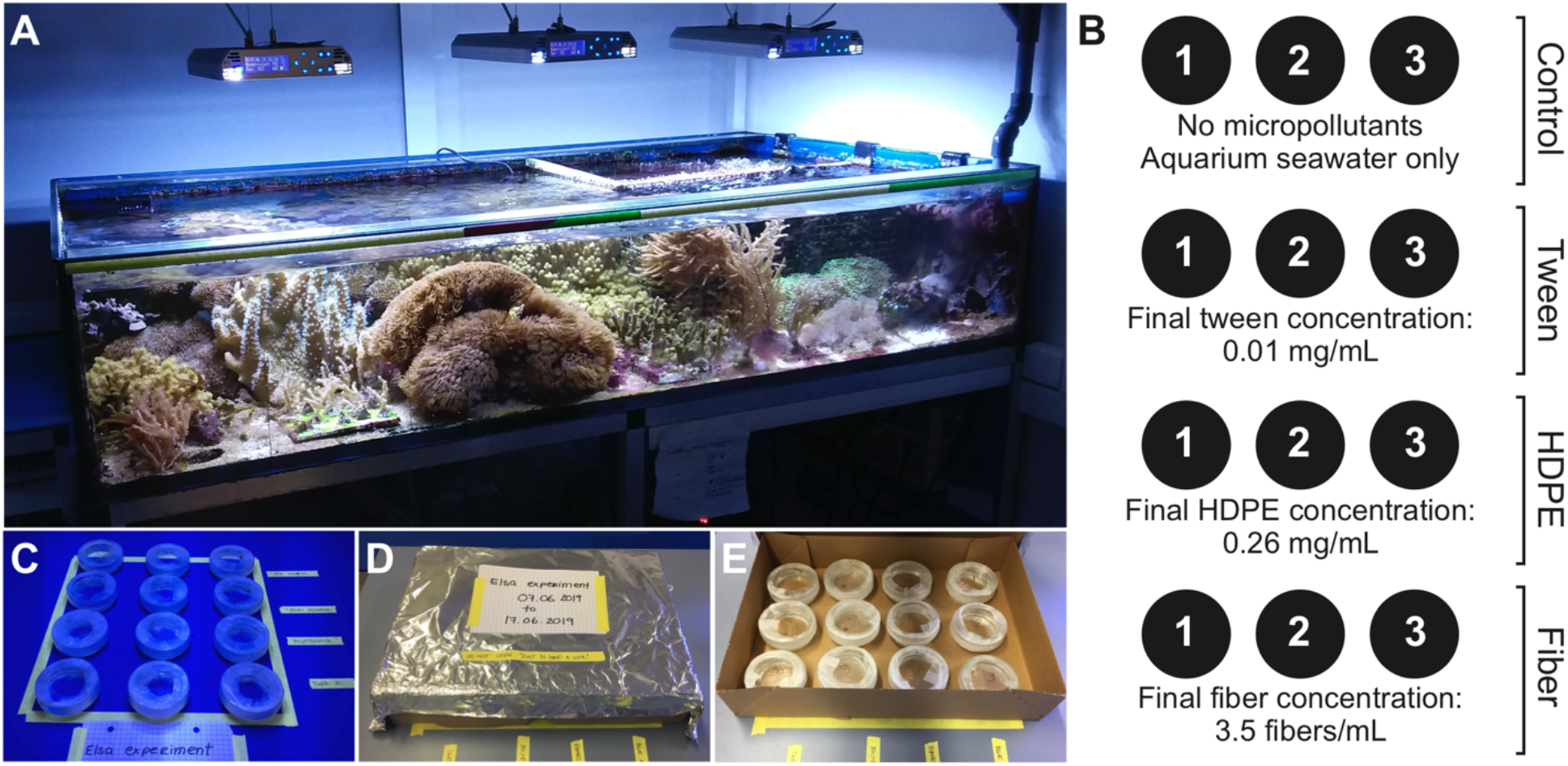
Experimental setup. A) Aquarium hosting a small reef ecosystem from which 15 mL was transferred into each incubation petri dish. B) incubation set of twelve glass petri dishes and associated incubations. C) Experimental display under artificial sunlight. D) and E) experimental display with limited access to light.

The oxygen concentration in each incubation was closely monitored over the first 48 h of the experiment using a Stand-alone Fiber Optic Oxygen Meter (PreSens Fibox 4) interacting with the oxygen sensor spots. At the beginning of the experiment, four samples of 15 mL were collected from the aquarium to assess the initial microbial community (T0; referred to as “aquarium”). All incubations and aquarium samples were processed through 4 or 15 mL Amicon® Ultra Centrifugal Filters (4000 rpm, RCF 3399 *g, 10 min. at 20 °C) to concentrate microbes and associated particles depending on the incubation. The concentrated supernatant was equally transferred in two Lysing matrix E tubes for every incubation.

### Quantitative PCR

The DNA was extracted as in Pichler *et al.* (2018), using 1 mL of a C1 extraction buffer ^14^. To lyse cells, the samples were subsequently heated at 99 °C for 2 min, frozen at −20 °C for 1 h, thawed at room temperature, and heated again at 99 °C for 2 min. After homogenization and centrifugation, the samples were concentrated a second time through Amicon filters, down to a final volume of ca. 100 *μ*L. The supernatant was purified following the DNeasy® PowerClean® Pro Cleanup Kit (Qiagen, Hilden, Germany). To assess the number of 16S copies at the end of the experiment, all samples were amplified using quantitative PCR (qPCR; Bio-Rad CFX connect™ Real-Time System). Every reaction contained 4*μ*L of DNA template, 10 *μ*L of Supermix, 5.2 *μ*L H_2_O, and 0.4 *μ*L of forward and reverse primer, and were subject to the following PCR program: denaturation at 95 °C for 3 min, and 40 amplification cycles (denaturation at 95 °C for 10 s, annealing 55 °C for 30 s). All qPCR reactions were set up using an Eppendorf EpMotion pipetting robot that has <5% technical variation and results in qPCR reaction efficiencies (standard curves) having >90% ^15^.

### 16S amplicon library preparation

To assess the diversity of the microbial community in the experimental samples, the V4 hypervariable region 16S rRNA gene (ca. 250 base pairs) was amplified with a set of primers (515F 5’-TATGGTAATTGTGTGCCAGCMGCCGCGGTAA-3’ and 806R 5’-AGTCAGTCAGCCGGACTACHVGGGTWTCTAAT-3’), combined to a forward (P5) and reverse (P7) adaptor, and unique dual indices for every sample ^14^. The preparation of the samples for the polymerase chain reaction (PCR) was done according to Pichler *et al.* (2018). In short, 4 *μ*L of extracted DNA was mixed to 5 *μ*L 5× PCR buffer, 1 *μ*L 50 mM dNTP, 1 *μ*L forward 515F and 1 *μ*L reverse 806R primers, 9.9 *μ*L H_2_O, 3 *μ*L MgCl_2_ and 0.1 *μ*L Taq DNA polymerase, for a total volume of 25 *μ*L for each sample. The amplification took place under specific PCR settings: denaturation at 95 °C for 3 min, 35 amplification cycles (denaturation at 95 °C for 10 s, annealing 55 °C for 30 s, elongation 72 °C for 1 min), and elongation at 72 °C for 5 min to ensure polymerization of all amplified DNA strands. PCR products were run through a 1.5% (w/v) agarose gel, and DNA strands were subsequently extracted using the QIAquick® Gel Extraction Kit (Qiagen, Hilden, Germany). The DNA concentration was quantified using the fluorometer QuBit 2.0 (Life Technologies, Grand Island, USA) and its associated dsDNA high-sensitivity assay kit. As preparation for 16S amplicon sequencing, all samples were pooled together by adding 5 *μ*L of every sample at a DNA concentration of 1 nM.

### 16S amplicon sequencing

A high diversity library was added to the 16S amplicon pool to enhance the recognition of the 16S sequences by the Illumina MiniSeq. The DNA was denatured by adding of 0.1 nM NaOH for a short period of 5 min, which was then directly neutralized with a tris-HCl buffer (pH 7) to avoid hydrolyzation of the DNA. To not overload the flow cell, a two-step dilution was performed on the 16S pool for a final DNA concentration of 1.8 pM, resulting in a final volume of 500 *μ*L MiniSeq solution. Four additional sequencing primers after ^14^ were added to successfully undergo the dual-index barcoding with the MiniSeq. Finally, the prepared 1.8 pM solution of 16S, transcriptomes, and the four sequencing primers was loaded into the reagent cartridge.

### Data analysis

To transform the demultiplexed sequences from Illumina MiniSeq into an OTU table, the raw data was manipulated using USEARCH v11.0 (https://drive5.com/usearch) ^16^ following the method developed by Pichler and colleagues (2018). Most similar sequences sharing at least 97% of bases were grouped, and associated to an operational taxonomic unit (OTU). Each OTU was classified within the Taxonomic Classification System using MacQiime v1.9.1 (http://qiime.org/). To keep a control on the analyzed data, OTUs of less than 10 reads to all samples were discarded. Statistical analyses were computed in R v3.3.3 ^17^. For phylogenetic reconstruction, most abundant selected OTUs were identified using blastn (BLAST®, https://blast.ncbi.nlm.nih.gov/). Sequences were aligned in MAFFT v7.427 (https://mafft.cbrc.jp/alignment/software/). The phylogenetic tree was inferred using Seaview v4.7 ^18^ under PhyML optimized settings (GTR model), including 100 bootstrap replicates ^19^. All related primary data and R scripts are stored on GitHub (https://github.com/PalMuc/PlasticsBacteria).

### Raman spectroscopy

Forty textile fibers were randomly subsampled and their associated spectrum obtained with a HORIBA JOBIN YVON XploRa ONE micro Raman spectrometer belonging to the Mineralogical State Collection Munich (SNSB). The used Raman spectrometer is equipped with edge filters, a Peltier cooled CCD detector and three different lasers working at 532 nm (green), 638 nm (red) and 785 nm (near IR). To perform the measurements the near IR Laser (785 nm) was used, with a long working distance objective (LWD), magnification 100× (Olympus, series LMPlanFL N), resulting in a 0.9 *μ*m laser spot size on the sample surface. The wavelength calibration of the IR laser was performed by manual calibration with a pure Si wafer chip, the main peak intensity had values in the interval 520 cm^−1^ +/− 1 cm^−1^. The wave number reproducibility was checked several times a day providing deviation of less than < 0.2 cm^−1^. Monthly deviation was in the range of 1 cm^−1^ before calibration. The necessary power to obtain a good-quality spectrum varied between 10% and 50% (i.e., respectively 2.98 mW and 18 mW +/− 0.1 mW on the sample surface) depending on the type and degraded stage of the measured textile fiber. The pin-hole and the slit were respectively set at 300 and 100. Each acquisition included two accumulations with a grading of 1200 T and an integration time of 5 s over a spectral range of 100 to 1600 cm^−1^. The precision of determining Raman peak positions by this method is estimated to be ± 1 to ± 1.5 cm^−1^. Resulting Raman spectra were analyzed using LabSpec Spectroscopy Suite software v5.93.20, treated in R v3.3.3, manually sorted in Adobe Illustrator CS3, and compared with available spectra from published work. All related Raman spectra and R scripts are stored on GitHub (https://github.com/PalMuc/PlasticsBacteria).

## RESULTS & DISCUSSION

For four days, a coral reef aquarium microbial community was incubated with Tween 20, HDPE microbeads and intertidal textile fibers in a microcosm experiment to test its potential to bioremediate widespread micropollutants. After sequencing of the V4 hypervariable region of the 16S rDNA genes a total of 1,463,028 sequences were obtained, from a total of 50 samples. After the quality control on the data, all sequences were clustered in 3,884 OTUs, of which 563 (85%) could be taxonomically classified.

### Respiration and induced microbial activity

Ten to 12 hours after the beginning of the experiment, a noticeable decrease in oxygen concentration was measured in all incubations containing micropollutants that was not observed in the control (Fig. 2A). This increased oxygen consumption in the presence of micropollutants indicates that microbial metabolism was stimulated by these pollutants, and their utilization as a carbon source for growth. It is likely that the transition between initial and final microbial communities initiated at this time in all incubations, which correlates with the theory that plastic surfaces are colonized within 24 h in the natural environment ^6^.

**Figure 2.**
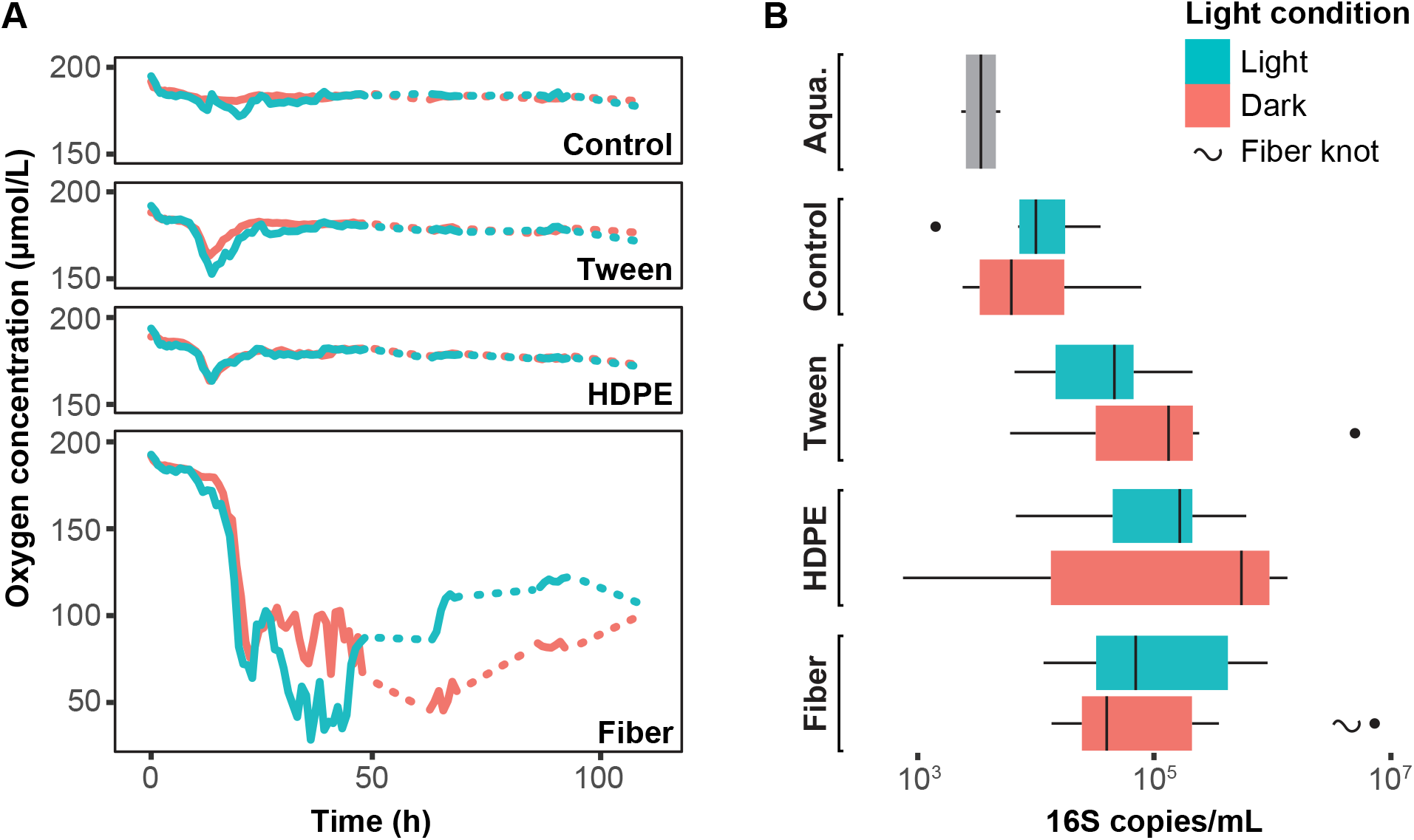
A) Oxygen respiration measured in the four incubations (average of three replicates. Dashed lines represent missing data (nights where no measurements were taken). B) 16S rRNA gene copies measured with qPCR in the same four incubations at the end of the experiment. Error bars represent standard deviations across three biological replicate incubations.

Control, tween and HDPE incubations reached an equilibrium between oxygen consumption and production after this time, whereas microbial communities in the textile fiber incubations continued to consume oxygen at a high rate (Fig. 2A). This indicates a higher microbial activity induced by the intertidal textile fibers. These fibers were mainly pigmented with a black dye (CI. reactive black 5 or 8) and a blue dye (phthalocyanine 15 (PB15)) according to the Raman analysis ^20,21^. Indeed, as much as 42% of the fiber spectra expressed only the fiber pigment, covering the fabric signal and preventing the identification of the polymer composition (Fig. 3). Here, only cotton expressed a signal strong enough to be recognized in the Raman spectra, with characteristic Raman peaks at positions 379, 435, 953, 1091 and 1116 cm^−1^ (Fig. 3). Hence, at least 44% of all fibers collected in the intertidal zone were identified as cotton, which appears to be one of the most abundant fiber materials found in the environment ^11,22^.

**Figure 3.**
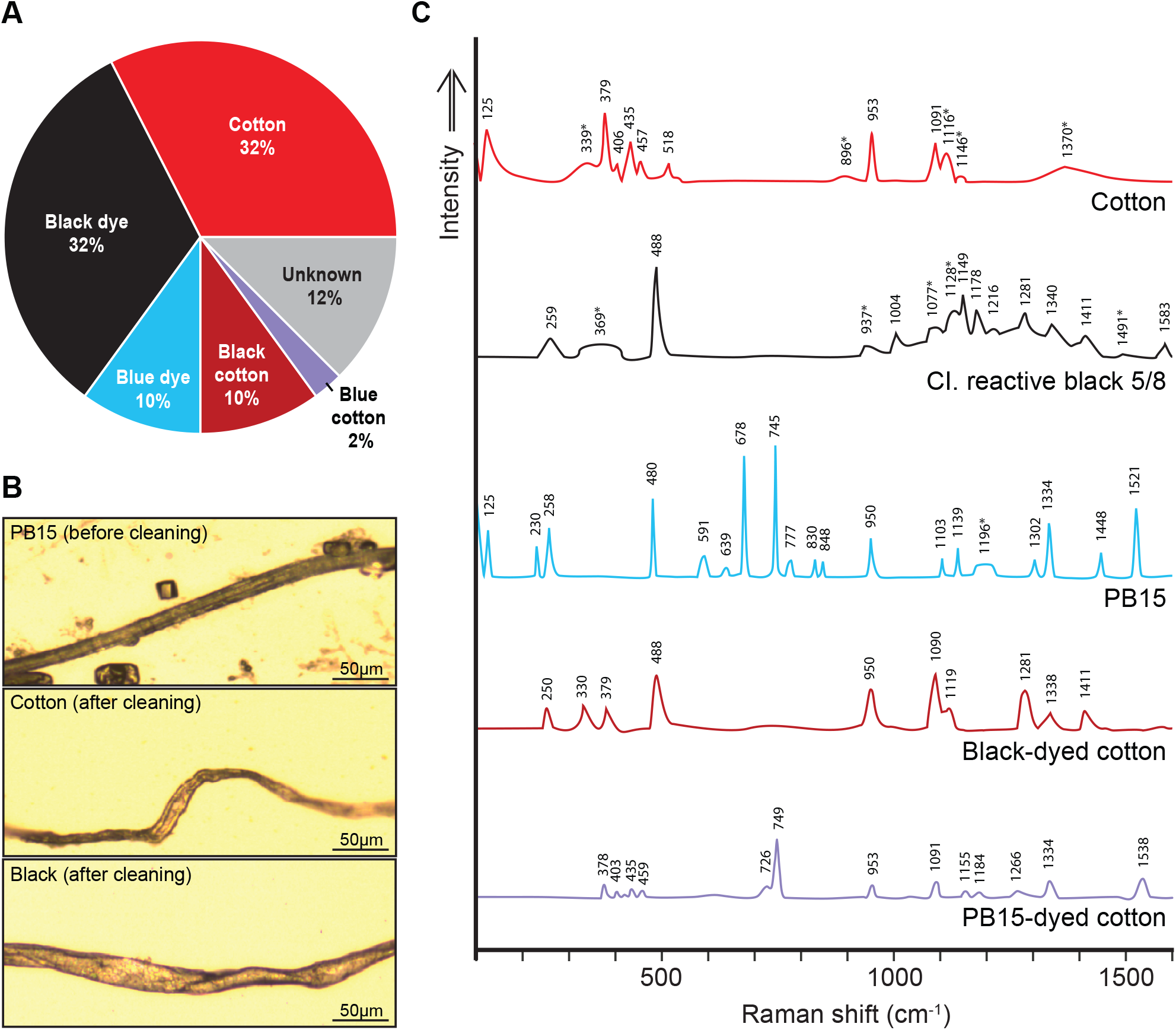
Analysis of the textile fiber sample extracted from the intertidal sediment at Coral Eye Resort. A) fiber type ratio based on signals obtained with Raman spectroscopy. B) photograms of different fibers measured with Raman spectroscopy, illustrating examples of fibers before and after cleaning. C) Raman spectra of five fiber types identified. Numbers indicate the position of the peaks (cm^−1^) and the associated asterisk (*) indicates a broad peak. Note: in 42% of the measurements, only the pigment signals were expressed, covering the polymer signature and preventing the identification of the fabric of those fibers. Only cotton seems to have a signal strong enough to overcome the pigment signature.

As textile fibers were sampled directly from an intertidal sandy beach in Indonesia (see Table 1), they were already exposed to high ultraviolet (UV) radiation and temperature, which are the main factors participating in polymer degradation and fragmentation on beaches ^23–25^. Indeed, UV radiation causes photooxidative degradation, which results in breaking of the polymer chains, produces free radicals and reduces the molecular weight, causing deterioration of the material, after an unpredictable time ^26^. This process rendered the textile fibers more subject to colonization in comparison to virgin HDPE microbeads, due to their advanced deteriorated stage, and likely facilitated the hydrolysis of carbon by hydrocarbon-degrading bacteria ^27^. Another study, by Romera-Castillo et al., demonstrated that irradiated plastic debris stimulated microbial activity in a mesocosm experiment in comparison to virgin plastics, supporting the results obtained in our study ^28^.

### Microbial community development

Another indication that microbial communities developed according to the provided carbon source (i.e., tween, HDPE microbeads and intertidal textile fibers) is the higher amount of 16S copies in tween, HDPE and textile fiber incubations measured using qPCR, in comparison to the control incubations (Fig. 2B). Moreover, microbial communities were significantly different (Analysis of Similarity: P < 0.01) between incubation types (Fig. 4), in comparison to the initial microbial community measure from the aquarium, which was dominated by the families Bacillaceae, Planctomycetaceae, Bacteriovoracaceae and Cellvibrionaceae (Fig. 5).

**Figure 4.**
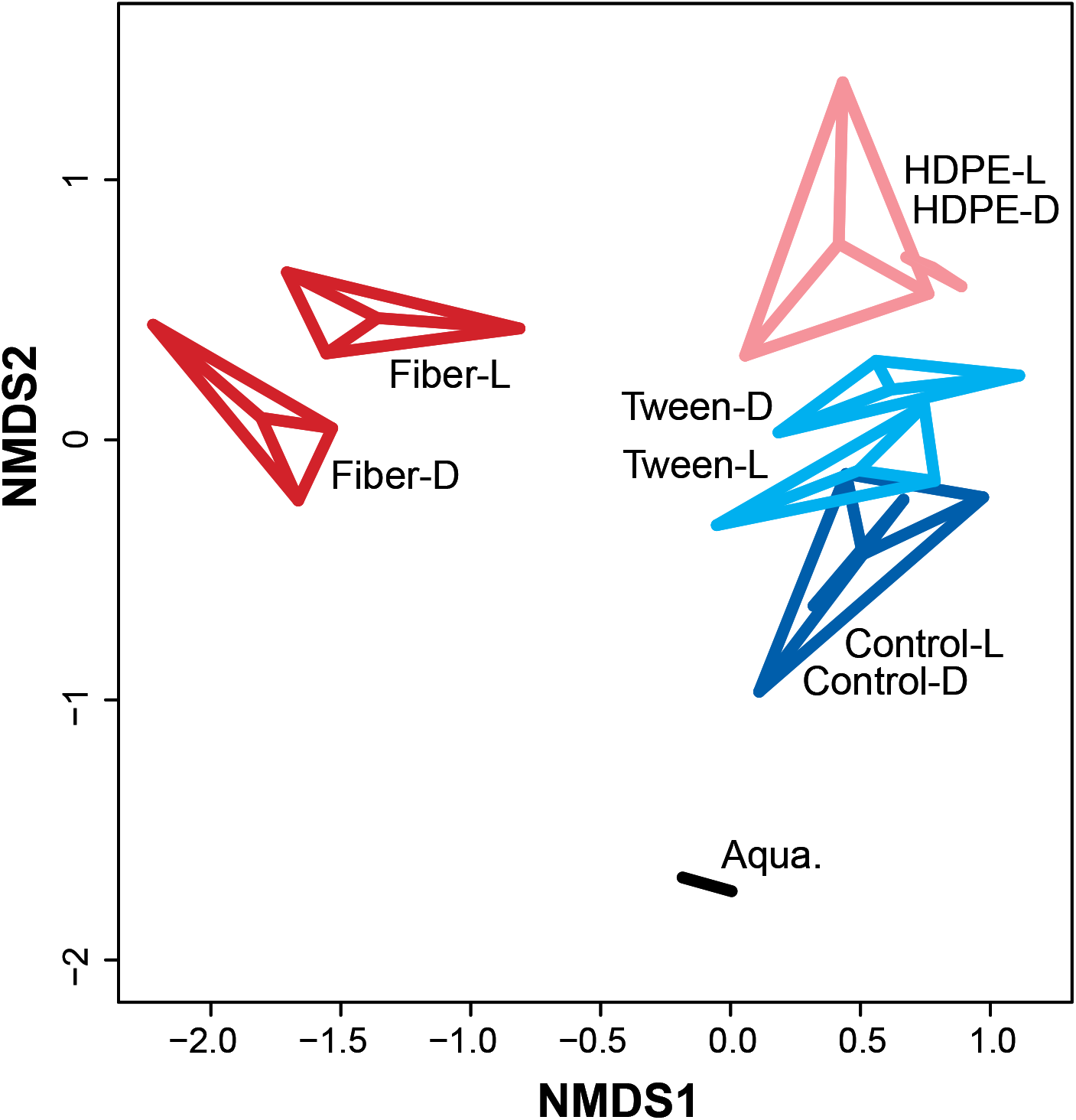
Non-metric multidimensional scaling (NMDS) analysis highlighting the difference in arbitrary distances between incubation-specific microbial communities (ANOSIM p = 0.001).

**Figure 5.**
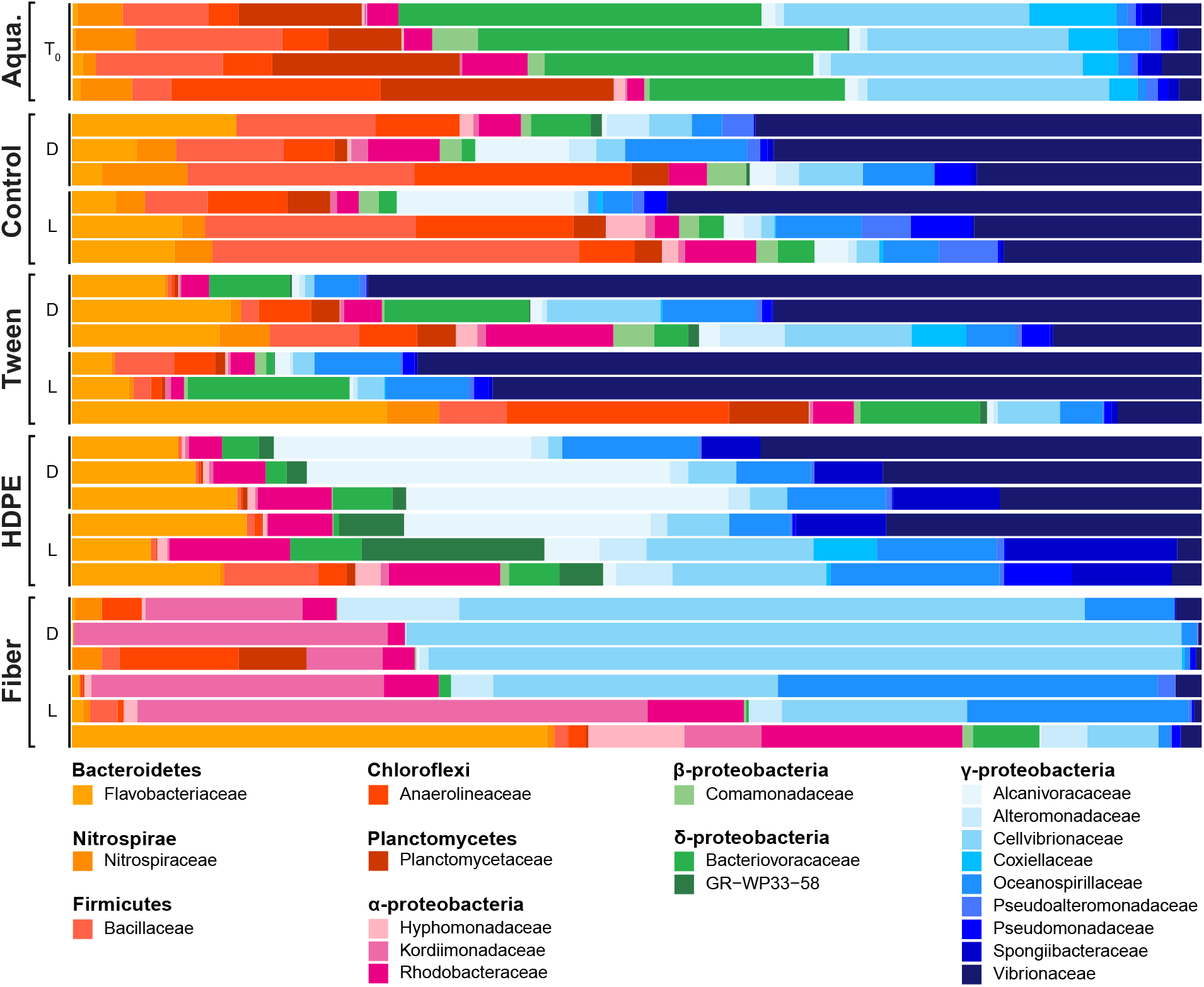
The relative proportion of the 20 most abundant families across all incubations, separated into incubation types and light conditions (T_0_: initial community; D: dark; L: light).

Flavobacteriaceae and Vibrionaceae were particularly ubiquitous in control, tween and HDPE incubations. However, HDPE-incubated communities were enriched with bacteria from the Family Alcanivoracaceae, whereas communities from the textile fiber incubations were enriched with bacteria from the Kordiimonadaceae and Cellvibrionaceae (Fig. 5). The observed difference between microbial communities from micropollutant-bearing and control incubations might also be explained by an accumulation of plastic-associated dissolved organic carbon (DOC) in the microcosms ^28^. Romera-Castillo et al. discovered that plastic debris releases a non-negligible amount of DOC, most of it leaches within the first few days after the initial contact with seawater, which would be applicable to the virgin HDPE microbeads ^28^.

Control and tween incubations had a highly variable microbial communities between replicates, whereas HDPE and textile fiber incubations showed very similar replicates. Nonetheless, all had differences in the community composition between light and dark settings (Fig. 4, 5). The disparity between microbial communities from incubations with HDPE microbeads and textile fibers indicates the development of polymer-dependent taxa, which is also supported by the findings of Frère et al. (2018). The families Oceanospirillaceae, Vibrionaceae, Flavobacteriaceae and Rhodobacteraceae are putative to the plastisphere with common representatives in HDPE and textile fiber incubations, however less diverse than previously observed in other studies ^3,5^. This might be related to the short experimental time; micropollutants and debris are otherwise accumulating over months and years in the ocean ^30^.

Operational taxonomic units (OTUs) of bacteria that were enriched in particular incubations were identified (Fig. 6). Thirty-one OTUs are shared between all incubation types and the aquarium water itself, of which two OTUs, i.e., OTU001 (γ-proteobacteria) and OTU004 (Flavobacteriia), were highly abundant across all incubations (Fig. 6). Only one taxon was especially enriched in the tween incubation and shared some OTUs with HDPE and control incubations, which were largely affiliated with the γ-proteobacteria. Textile fibers were especially colonized by α- and γ-proteobacteria, identified as first colonizers ^6,7^. HDPE incubations were mainly characterized by the development of Bacteroidetes (i.e., Family Flavobacteriaceae), additionally to α- and γ-proteobacteria, hypothesized to colonize plastics at a later stage ^6,7^.

**Figure 6.**
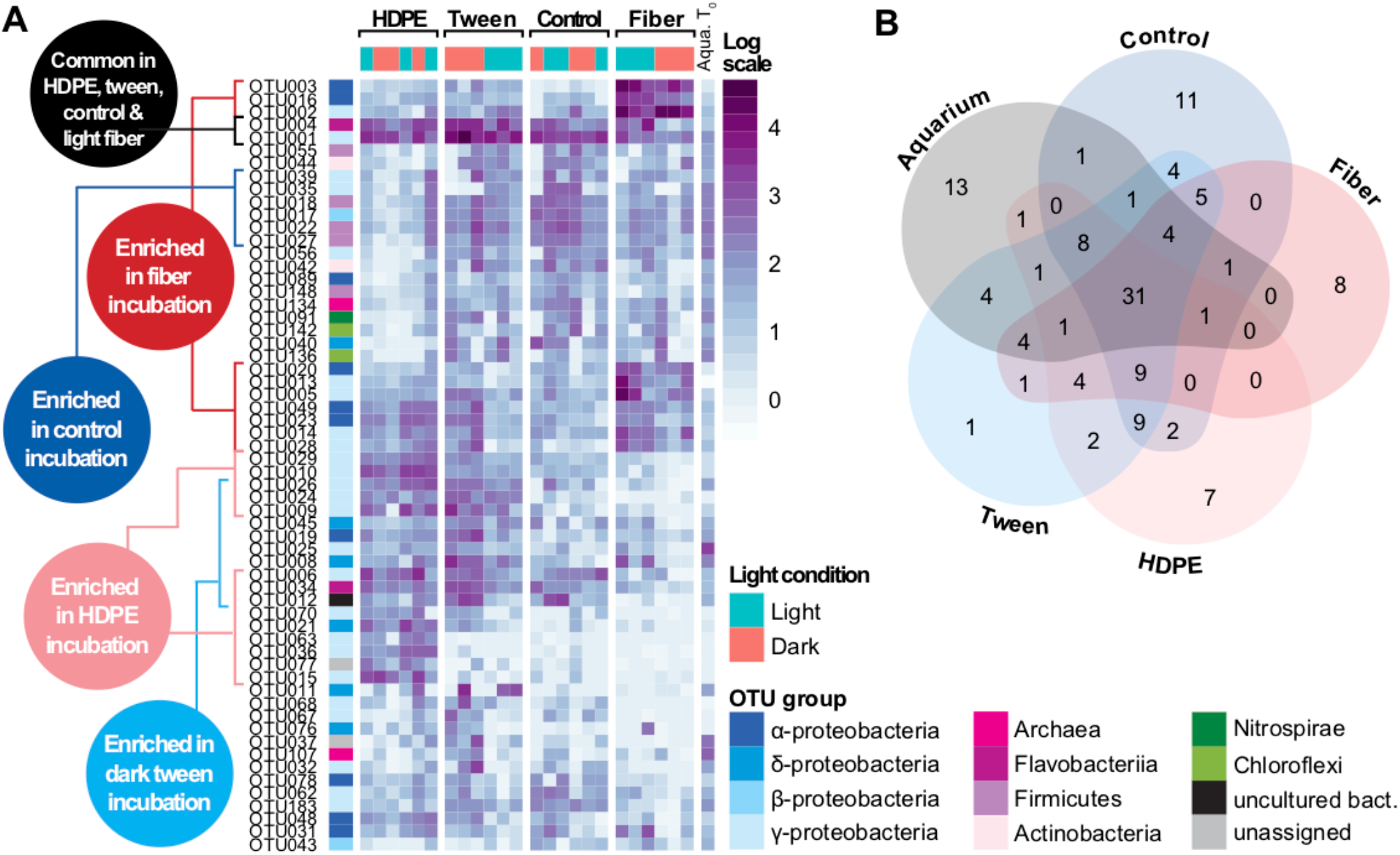
A) Log-scaled heat map highlighting the abundance of the 10 most abundant OTUs in each incubation. B) Venn diagram showing a core community, and incubation-specific OTUs.

### Impact of light on microbial development

The presence of light was correlated with the development of unique microbial communities, especially observable in HDPE and textile fiber incubations, similar to the findings of Fuhrman et al. (2008) and Sánchez et al. (2017) regarding the impact of light on the microbial communities ^31,32^. Indeed, the dark-incubated HDPE microbial community was dominated by the three families Vibrionaceae, Alcanivoracaceae and Flavobacteriaceae, whereas the light-grown HDPE microbial community was more diverse with a shared dominance between eight families (Rhodobacteraceae, GR-WP33-58, Spongiibacteraceae, Vibrionaceae, Cellvibrionaceae, Oceanospirillaceae, Alcanivoracaceae, and Flavobacteriaceae) (Fig. 5).

Some taxa were enriched under artificial sunlight when incubated with HDPE and textile fibers, but enriched in the presence of Tween 20 under dark conditions. This observation is well represented by OTU028 and OTU049, which were most abundant in textile fiber and HDPE incubations in the presence of light, otherwise dominant in the tween incubation under dark conditions. In general, OTUs enriched in Tween-20 incubations had a higher relative abundance in dark settings, also supported by our qPCR results. The flexibility these taxa show in using carbon from various sources (HDPE microbeads *vs* Tween 20) depending on the light availability might highlight an opportunistic behaviour ^33^. Another hypothesis suggests a higher level of available DOC in light-exposed textile fiber and HDPE incubations, caused by the polymer exposure to artificial sunlight ^13,28^. These findings may help us better understand the plastisphere dynamic in situations similar to, for example, microorganisms settled on plastic debris initially floating in the photic zone and later buried in the sediment or sinking in regions with limited light availability.

### Hydrocarbon-degrading bacteria

Several OTUs enriched in the different incubation types revealed to be closely related to known hydrocarbon degrading microorganisms (Fig. 7). This, together with the increased rates of oxygen consumption in those incubations, highlights their potential for biodegradation of organic carbon based micropollutants. The six most abundant OTUs of the textile fiber incubation (OTU002, −003, −005, −013, −016 and −020) are closely related to the genera *Kordiimonas* and *Defluviimonas* (α-proteobacteria), and *Simiduila*, *Marinobacterium* and *Neptuniibacter* (γ-proteobacteria) according to the inferred phylogenetic tree. OTU003 and −020 were also closely related to *K. gwangyangensis* (NR_043103.1), which can hydrolyze six different polycyclic aromatic hydrocarbons (PAHs) ^34^, giving this taxon a strong potential for microplastic bioremediation. The genus *Alteromonas*, hosting hydrocarbon-degrading microorganisms, has been previously identified as part of the plastisphere from North Adriatic Sea and Atlantic Ocean ^5,35^. More specifically, *Alteromonas naphthalenivorans* was identified as a naphthalene consumer ^36^. *Defluviimonas alba* was isolated from an oilfield water sample suggesting that it has a potential for degradation of hydrocarbon similarly to *Defluviimonas pyrenivorans* ^37^, however not tested for hydrocarbon hydrolysis ^38^. *Bowmanella pacifica* was identified during a search for pyrene-degrading bacteria ^39^, which is a PAH highly concentrated in certain plastics ^40^. *Bowmanella* spp. were also identified on polyethylene terephthalate (PET) specific assemblages from Northern European waters ^41^. Because these taxa were probably biofilm-forming microorganisms developed on the surface of textile fibers, it is very likely that they partly consumed the polymer, an available source of carbon and energy, owing to their capability to hydrolyze hydrocarbons. Although approximately half of the fibers used in the experiment were of natural fabric, it has been observed that cotton is a powerful sorbent used to treat oil spill ^42^. The latter suggest that cotton fibers may absorb traces of hydrocarbon present in the environment ^43^ and, therefore, provide a source of hydrocarbon for hydrocarbonoclastic bacteria.

**Figure 7.**
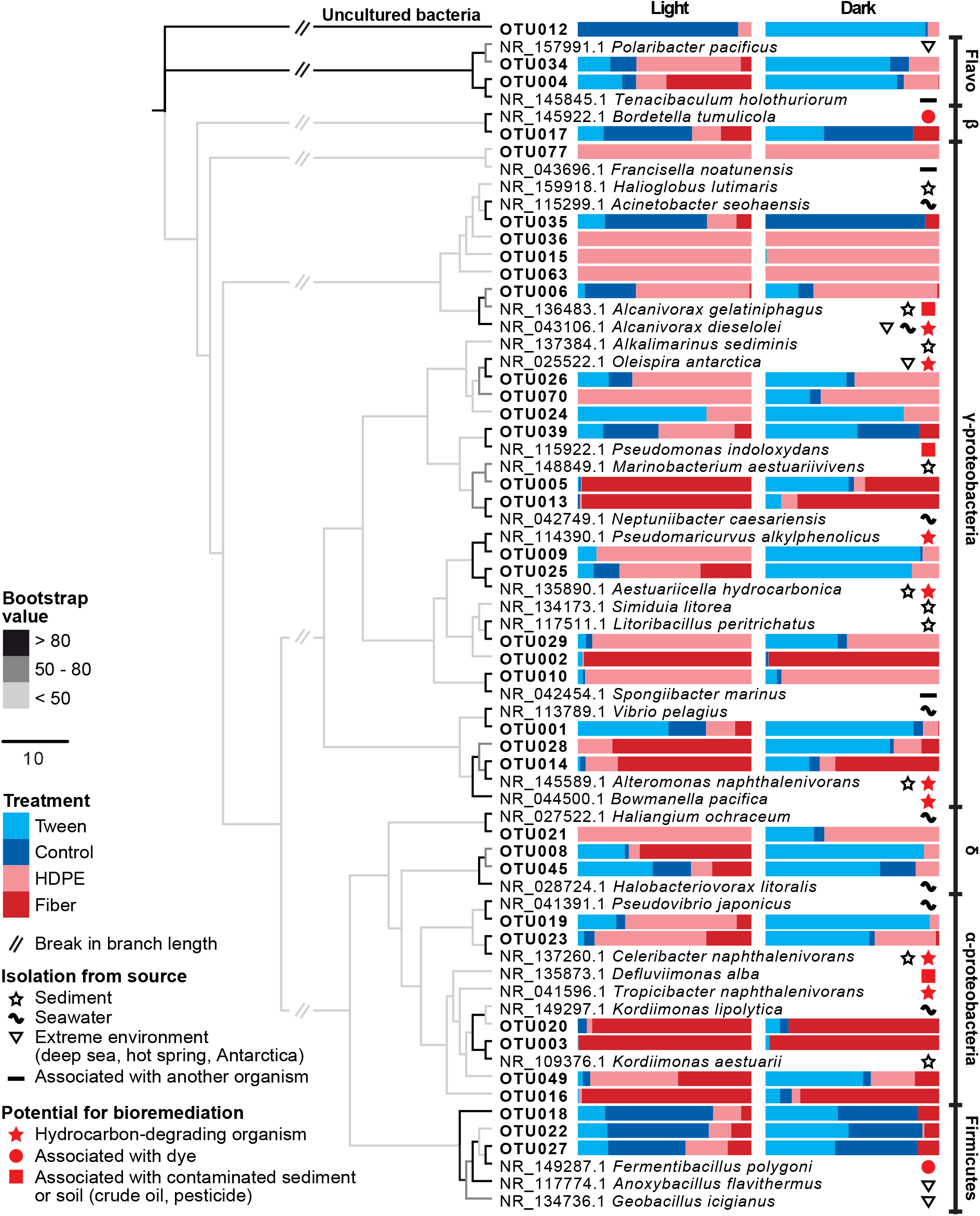
Phylogenetic reconstruction of 36 OTUs identified as incubation-enriched in Fig. 6 and their closest associated named species from NCBI database. The relative abundance of OTUs in each incubation (pale blue: tween; dark blue: control; pale red: HDPE; dark red: fiber) is represented by bar charts, divided according to the light condition. Named-species microorganisms were classified based on their isolation from the source (black star, wave, triangle and line) and marked for their potential to bioremediate microplastics (red star, circle and square).

The HDPE incubation stimulated the activity and enrichment of six OTUs (OTU001, −004, −006, −010, −015 and −026) closely related to hydrocarbon-degrading bacteria, different from those enriched in textile fiber incubations. For instance, OTU006 was affiliated with the genus *Alcanivorax*, which are specialized in degrading alkanes, especially in contaminated marine environments ^44^. This genus was also identified as being potentially important for PET degradation in the natural marine environment ^41^. All other five OTUs were abundant only in light-grown HDPE communities, which were affiliated with the taxa *Pseudomaricurvus alkylphenolicus*, *Celeribacter naphthalenivorans*, *Oleispira antarctica, Tropicibacter naphthalenivorans*, and *Aestuariicella hydrocarbonica*. Given their enrichment in the HDPE incubations in the presence of light, and the increased oxygen consumption relative to the control, it seems likely that these conditions selected for these specific organisms that were uniquely adept to utilize organic carbon from the HDPE as a carbon and energy source. This indicates their potential for the bioremediation of this plastic type. The different respiration rates of HPDE treatment and community assembly compared to the Tween 20 treatment (an emulsifier for the HDPE microbeads) and demonstrates a unique effect on microbial community formation due to the HDPE itself (as opposed to microbes that may just be eating the Tween that is coated on the HDPE).

### Extrapolating from our experiment to the natural environment

Although this study did not use “natural” microbial communities from the intertidal habitats from which the textile fibers derive, it tests for the potential response of microbial communities to widespread micropollutants. Since the identified hydrocarbonoclastic bacteria in our incubations are also common to both natural marine sediments and water column ^5,34,39,41,44^, this study contributes to the developing understanding of how hydrocarbon-degrading bacteria utilize various fabric types as a carbon and energy sources during bioremediation processes. Three main lines of evidence indicate that the plastics and textile fibers were consumed by hydrocarbon-degrading microorganisms during the experiment: (1) higher rates of oxygen consumption relative to the control, (2) increased abundance as indicated by qPCR and (3) the development of unique microbial communities that form in incubations containing different types of micropollutants. Moreover, the same groups of hydrocarbonoclastic bacteria found in our study are present in the marine environment. The findings resulting from this study also demonstrated that light availability is an important factor shaping hydrocarbon-degrading bacterial communities. We speculate that this is due to photochemical dissolution of the plastic and textile substrates, which might allow for more readily bioavailable substrates for the developing biofilms. A deeper study into the topic is necessary to enlarge our understanding behind microbial adaptation to changing environmental conditions, including photochemical dissolution of plastic waste.

## ACKNOWLEDGEMENTS

This study was conducted within the frame of the program Lehre@LMU, part of the Geobiology and Paleobiology section of the Department of Earth and Environmental Sciences, Ludwig-Maximilians-Universität München. We thank Coral Eye Resort (Marco Segre Reinach) for providing the textile fibers and the Indonesian authorities for providing the research visa and permit (research permit holder: Elsa Girard; SIP no.: 97/E5/E5.4/SIP/2019). We are also grateful for the time Aurèle Vuillemin, Ömer Coskun, Paula R. Ramirez, and Nicola Conci took to help us in the lab and with the data analysis.

This study was funded by Lehre@LMU (project number: S19_F2.; Studi_Forscht@GEO), and to budget funds to WO and GW. GW acknowledges support by LMU Munich’s Institutional Strategy LMUexcellent within the framework of the German Excellence Initiative for aquarium set-up and maintenance.

